# Novel insights into the morphology of *Plesiochelys bigleri* from the early Kimmeridgian of Northwestern Switzerland

**DOI:** 10.1101/582700

**Authors:** Irena Raselli, Jérémy Anquetin

## Abstract

Plesiochelyidae were relatively large coastal marine turtles, which inhabited the epicontinental seas of Western Europe during the Late Jurassic. Their fossil record can be tracked in Germany, Switzerland, the United Kingdom, France, Spain and Portugal. The Jura Mountains, in northwestern Switzerland, have been the main source for the study of this group, mostly thanks to the rich and famous historical locality of Solothurn. In the last two decades, numerous plesiochelyid remains have been collected from Kimmeridgian deposits (Lower *Virgula* Marls and Banné Marls) in the area of Porrentruy (Canton of Jura, Switzerland). This material was revealed by construction works of the A16 Transjurane highway between 2000 and 2011, and led to the recent description of the new species *Plesiochelys bigleri*. In the years 2014 and 2016, new fragmentary turtle material was collected from the Banné Marls (Reuchenette Formation, lower Kimmeridgian) near the village of Glovelier, Canton of Jura, Switzerland. The new material consists of a complete shell, additional shell elements, a few bones from the appendicular and vertebral skeleton, and a fragmentary basicranium. This material can be confidently assigned to the species *P. bigleri*. It supports the presence of this species in the Banné Marls, slightly extends its spatial distribution and confirms the differences with the closely related species *P. etalloni.* The new material reveals that the split between the cerebral and palatine branches of the internal carotid artery occurs in a vertical plane in *P. bigleri*. This condition could not be observed in the type material due to poor preservation. This new character clearly distinguishes *P. bigleri* from *P. etalloni* and seems to be unique among thalassochelydians.

## Introduction

Thalassochelydia is a clade of coastal marine turtles that diversified mostly in Europe during the Late Jurassic (for a review, see [1]). Some thalassochelydians, namely the Eurysternidae Dollo, 1886 [2], were of small to medium size and lived primarily in lagoons and near-shore environments. Other thalassochelydians, such as the Plesiochelyidae Baur, 1888 [3] and Thalassemydidae Zittel 1889 [4], reached larger sizes (over 40 cm in carapace length) and were thought to thrive in more open marine conditions. However, the absence of stiffened paddles suggests that these turtles remained relatively close to the coast and were not able to cross large oceanic expanses [1]. Historically, the most productive localities for thalassochelydians were Solothurn in Switzerland and several plattenkalk localities in France and southern Germany (see [1] and references therein). In the past 20 years, extensive excavations have been carried out in northwestern Switzerland for the construction of the A16 Transjurane highway and have led to the discovery of several vertebrate-bearing layers in the Kimmeridgian of the region of Porrentruy, Canton Jura, Switzerland [5]. Two stratigraphical layers were particularly productive for vertebrates: the upper Kimmeridgian Lower *Virgula* Marls and the lower Kimmeridgian Banné Marls [6,7]. Thalassochelydians were among the most common vertebrates with around a hundred of more or less complete shells, a few skulls and several thousands of isolated remains [8]. The turtle material from the Kimmeridgian of the Porrentruy region was recently described in a series of papers [9–12]. Most of this material is referable to two closely-related species: *Plesiochelys etalloni* and *Plesiochelys bigleri* [11].

In the present paper, we describe new turtle material from the lower Kimmeridgian Banné Marls in Glovelier, Canton Jura, Switzerland, including a basicranium and a near complete shell. This material can be confidently referred to *Plesiochelys bigleri* and provides new important information on this recently described species.

## Material and Methods

### Material

The material presented herein was found in two times. A first limestone block was collected in 2014 by the first author. It contained the fragmentary remains of a shell (some costals and two peripherals), a partial scapula and a fragment of the basicranium. Two years later, new turtle material was recovered at the exact same spot in the outcrop. This turned out to be a near complete shell with some elements of the appendicular and axial skeleton (see below), and a poorly preserved fragment of a costal. One of the peripherals found in 2014 belongs to the near complete shell found in 2016 and was replaced during preparation. This confirms that all of this material was associated in the field.

MJSN CBE-0001 corresponds to most of the postcranial material collected in 2014, including an articulated fragment of the carapace (left costals 5–7 and peripherals 8 and 9), fragments of costals, an isolated peripheral, and a partial right scapula. MJSN CBE-0002 consists of a fragmentary basicranium and associated columella auris found in 2014. MJSN CBE-0003 corresponds to the near complete shell found in 2016. Associated with this shell, are a cervical and a caudal vertebrae, the left femur, the right pubis (still embedded in the matrix), and a poorly preserved fragment of a costal that evidently pertains to another individual.

The material discussed herein belongs at least to two individuals, represented by the articulated carapace fragment (MJSN CBE-0001) and the near complete shell (MJSN CBE-0003). The basicranium (MJSN CBE-0002) may belong to either of these individuals, or less likely represent a third one.

### Geological setting

The new material was collected from the Banné Marls at a locality called Combe du Bé (CBE; 47° 20’ 08’’ N, 7° 11’ 29’’ E) near the village of Glovelier, Canton of Jura, Switzerland (Fig. 1). The Banné Marls belong to the Reuchenette Formation and are dated from the Cymodoce ammonite zone (late early Kimmeridgian; Fig. 2; [7]), which makes them somewhat older than the Lower *Virgula* Marls (middle of the Reuchenette Formation, Eudoxus ammonite zone; [7]) and the famous Solothurn Turtle Limestone (top of the Reuchenette Formation, Autissiodorensis ammonite zone; [6,13]).

**Fig. 1.**
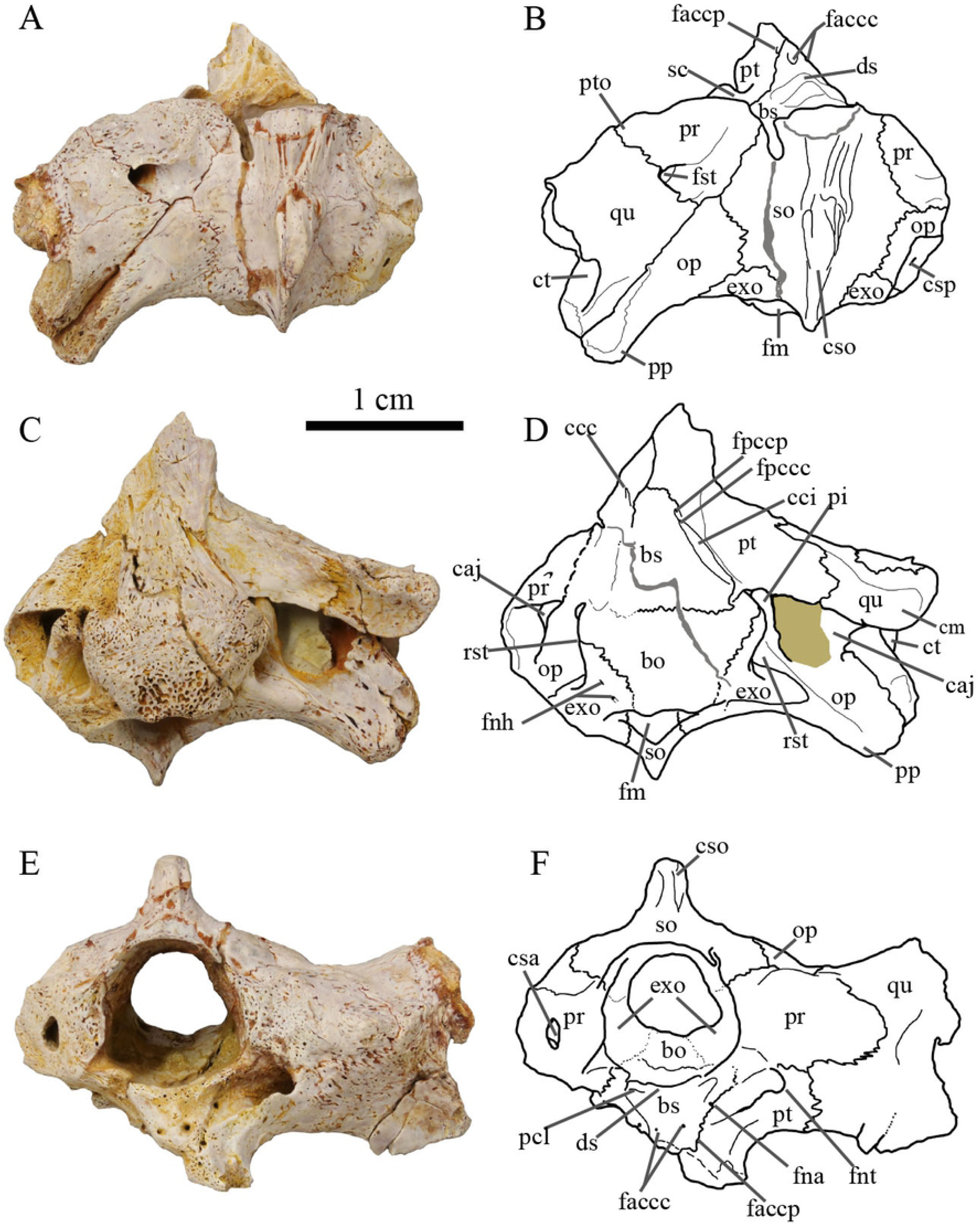
Map of the locality. The star marks the location of the outcrop in the small valley Combe du Bé near Glovelier, Canton of Jura, northwestern Switzerland.

**Fig. 2.**
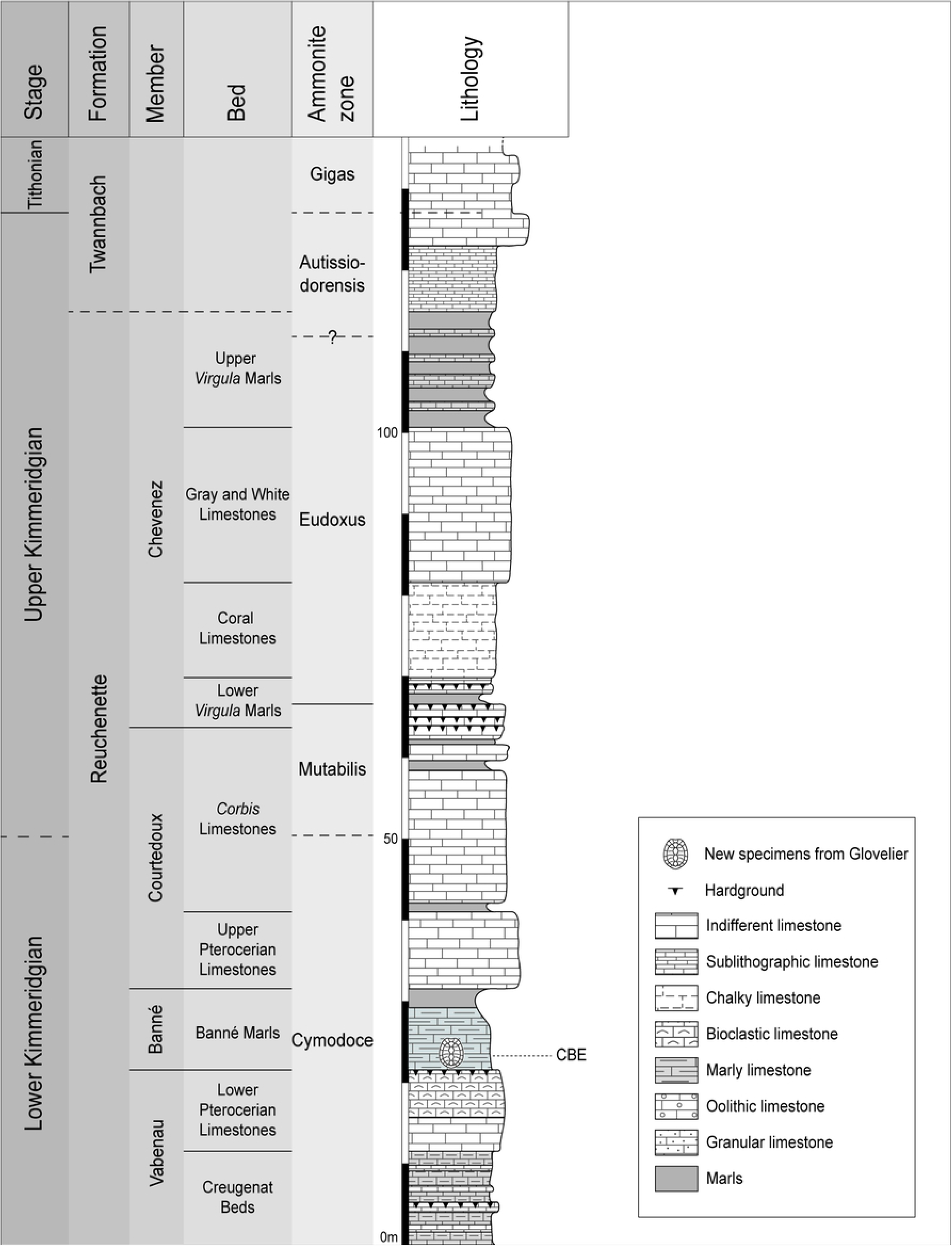
Stratigraphical column of the Reuchenette Formation in northwestern Switzerland. The shell marks the position of the new specimens in the Banné Marls (CBE: Combe du Bé, name of the locality). Modified from Püntener et al. [11].

The Banné Marls consist of fossiliferous marly limestones rich in bivalves, gastropods, brachiopods, echinoderms and cephalopods, as well as remains of vertebrates such as crocodylomorphs, turtles and fishes representing a diverse coastal marine biocenosis [7]. *Tropidemys langii* is the most common turtle taxon in this layer [9]. The new turtle material presented herein was found at the base of the Banné Marls, just above the hardground (Fig. 2).

### Comparative anatomy

The new material from Glovelier was primarily compared with the rich coeval Kimmeridgian turtle assemblage described recently from Courtedoux and Porrentruy, Canton of Jura, Switzerland (for an overview see [8] and references therein). The material referable to the species *Plesiochelys etalloni* and *Plesiochelys bigleri* was described in detail by Püntener et al. [11]. Comparisons also included recently revised material of *Plesiochelys* spp. from other parts of Switzerland and Europe [12,14–16].

First hand comparisons were made, notably for cranial anatomy, with the type material of *Plesiochelys bigleri* (MJSN TCH007-252 and MJSN TCH006-1451) and with beautifully preserved specimens of *Plesiochelys etalloni* from Solothurn (NMS 40870 and NMS 40871). The anatomical description of the cranium follows the nomenclature of Gaffney [17,18], updated by Rabi et al. [19]. The description of shell material follows the nomenclature of Zangerl [20].

### 3D models

#### Crania

The following crania were micro CT-scanned in order to gain insight into their internal anatomy and to look for potential differences of systematic value: MJSN CBE-0002, MJSN TCH007-252 (holotype of *Plesiochelys bigleri*), and NMS 40870 (*Plesiochelys etalloni*). The scans have been obtained at the University of Fribourg (Department of Geosciences). Scan specifications are listed in Table 1. The scans of specimens TCH007-252 (holotype of *Plesiochelys bigleri*), and NMS 40870 (*Plesiochelys etalloni*) were segmented with the 3D software Amira 6.0. The 3D models of the skulls and isolated bones can be accessed on MorphoSource (see data avaiability). The scan of the Glovelier specimen, MJSN CBE-0002, was not segmented due to insufficient contrast between the bone and the matrix.

**Table 1:** Scan specifications for the examined specimens.

| Specimen | Micro CT-Scanner | Voltage [kV] | Current [μA] | Filter [mm] | Rotation steps [°] | Voxel size [μm] |
| --- | --- | --- | --- | --- | --- | --- |
| MJSN CBE-0002 |  | 80 | 606 | Ti 0.5 | 0.2 | 25 |
| MJSN TCH007-252 | Bruker SkyScan<br>2211 | 190 | 50 | - | 0.2 | 39 |
| NMS 40870 |  | 190 | 59 | - | 0.2 | 53 |

#### Shell

The near complete shell MJSN CBE-0003 suffered only minor postmortem deformation and represents the best specimen to document the natural 3D shape of the shell of *Plesiochelys bigleri* (see below). In order to facilitate future comparisons, this shell was scanned with a structured-light surface scanner (Artec Space Spider, Artec Group), and a 3D model was computed with the software Artec Studio 13 Professional. The 3D model of the shell can be accessed on MorphoSource (seedata avaiability).

## Abbreviations

### Institutions

MJSN: Jurassica Museum (formerly Musée jurassien des sciences naturelles), Porrentruy, Switzerland
NMS: Naturmuseum Solothurn, Solothurn, Switzerland
NMBE: Naturhistorisches Museum Bern, Bern, Switzerland

### Localities

CBE: Combe du Bé, Glovelier, Switzerland
TCH: Tchâ foué, Courtedoux, Switzerland

## Systematic Paleontology

TESTUDINES Batsch, 1788 [21]

PAN-CRYPTODIRA Joyce, Parham & Gauthier, 2004 [22]

THALASSOCHELYDIA Anquetin, Püntener & Joyce, 2017 [1]

PLESIOCHELYIDAE Baur, 1888 [3]

### *Plesiochelys* Rütimeyer, 1873 [23]

Type species: *Plesiochelys solodurensis* Rütimeyer, 1873 [23]

### *Plesiochelys bigleri* Püntener, Anquetin & Billon-Bruyat, 2017 [11]

Type material: MJSN TCH007-252 (holotype), a disarticulated shell, consisting of the sub-complete carapace, the epiplastra, entoplastron, hypoplastra and left xiphiplastron, the posterior part of the cranium, and some elements of the appendicular skeleton [11]. MJSN TCH006-1451 (paratype), an isolated partial cranium [11].

Type locality and horizon: Tchâ foué (TCH), Courtedoux, close to Porrentruy, Canton of Jura, Switzerland. Lower *Virgula* Marls, Chevenez Member, Reuchenette Formation, late Kimmeridgian [6,7].

Occurrence: Early and late Kimmeridgian of the Porrentruy region, Canton of Jura, Switzerland [11]; early Kimmeridgian of Glovelier, Canton of Jura, Switzerland (this study).

Diagnosis: see [11] and [1].

Referred material: MJSN CBE-0001, shell fragments including articulated and isolated costals, a peripheral, and a partial scapula; MJSN CBE-0002, a partial basicranium with the left otic chamber and columella auris; MJSN CBE-0003, a near complete shell with the left femur, right pubis, a cervical vertebra, and a caudal vertebra.

## Description

The anatomy of *Plesiochelys bigleri* was recently described in detail [11]. The following description is therefore rather concise and focuses mainly on the new insights provided by the new material from Glovelier.

### Cranium

MJSN CBE-0002 consists of an incomplete basicranium and left otic chamber (Fig. 3). As preserved, the specimen is 31 mm in length and 40 mm in width. The medial half of a columella auris is preserved as well. The parietals, squamosals, postorbitals, quadratojugals, as well as all of the elements of the orbitonasal and palatal regions are missing. The lateral margin of the left otic chamber and basioccipital region are damaged.

**Fig. 3.**
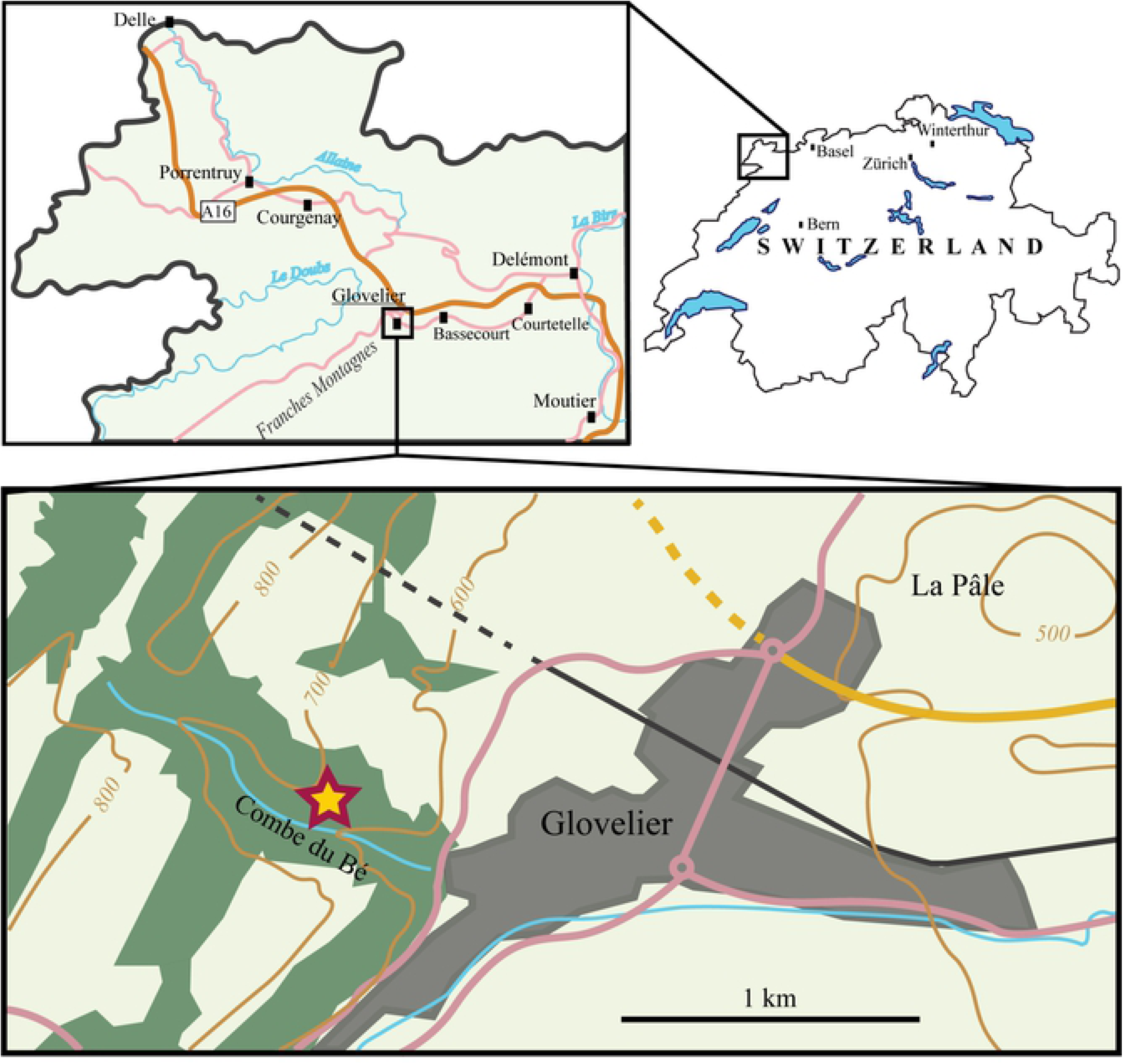
Basicranium MJSN CBE-0002. Photographs and interpretative drawings in dorsal view (A-B), in ventral view (C-D) and in anterior view (E-F). Abbreviations: bo, basioccipital; bs, basisphenoid; caj, cavum acustico-jugulare; ccc, canalis caroticus cerebralis; cci, canalis caroticus internus; cm, condylus mandibularis; csa, canalis semicircularis anterior; cso, crista supraoccipitalis; csp, canalis semicircularis posterior; ct, cavum tympani; ds, dorsum sellae; exo, exoccipital; faccc, foramen anterius canalis carotici cerebralis; faccp, foramen anterius canalis carotici palatinum; fm, foramen magnum; fna, foramen nervi abducentis; fnh, foramen nervi hypoglossi; fnt, foramen nervi trigemini; fpccc, formamen posterius canalis carotici cerebralis; fpccp, foramen posterius canalis carotici palatinum; fst, foramen stapedio-temporale; op, opisthotic; pcl, processus clinoideus; pi, processus interfenestralis; pp, processus paroccipitalis; pr, prootic; pt, pterygoid; pto, processus trochlearis oticum; qu, quadrate; rst, recessus scalae tympani; sc, sulcus cavernosus; so, supraoccipital.

The poor preservation of the lateral margin of the quadrate prevents direct comparison of MJSN CBE-0002 with the type material of *Plesiochelys bigleri*, which is remarkable in having a deep cavum tympani facing posterolaterally and a quadrate forming the complete anterior margin of the antrum postoticum [11]. Similarly, the processus articularis of the quadrate is damaged ventrally and the shape of the condylus mandibularis cannot be described. The imperfectly preserved processus trochlearis oticum is modest in development, differing from the condition in *Plesiochelys etalloni*.

As in the type specimens of *Plesiochelys bigleri*, MJSN CBE-0002 has a shallow pterygoid fossa and a reduced posterior extension of the pterygoid over the cavum acustico-jugulare (processus interfenestralis of the opisthotic largely visible in ventral view). This new specimen nicely documents the contacts of the pterygoid in the basioccipital region and allows us to complete the original description of *Plesiochelys bigleri*. The presence of a pterygoid-basioccipital contact is confirmed, as is the absence of contact between the pterygoid and exoccipital. Only the anteroventral part of the processus interfenestralis of the opisthotic has a sutural contact with the dorsal surface of the pterygoid. The pterygoid forms the posteroventral part of the foramen nervi trigemini, which appears to be rounded and relatively wide, in contrast to the slit-like opening found in *Plesiochelys etalloni* [11,12].

As preserved, the canalis caroticus internus is open ventrally. In contrast to the type specimens of *Plesiochelys bigleri*, both the foramen posterius canalis carotici cerabralis and the foramen posterius canalis carotici palatinum are clearly visible in MJSN CBE-0002. Interestingly, the palatine branch does not branch off laterally from the cerebral branch as usual in plesiochelyids and most turtles. Instead, the split between the cerebral and palatine branches occurs in a vertical plane and the foramen posterius canalis carotici palatinum is located anterior to the foramen posterius canalis carotici cerebralis (Fig. 4; see Discussion below). As far as we are aware of, this feature is unique to *Plesiochelys bigleri*.

**Fig. 4.**
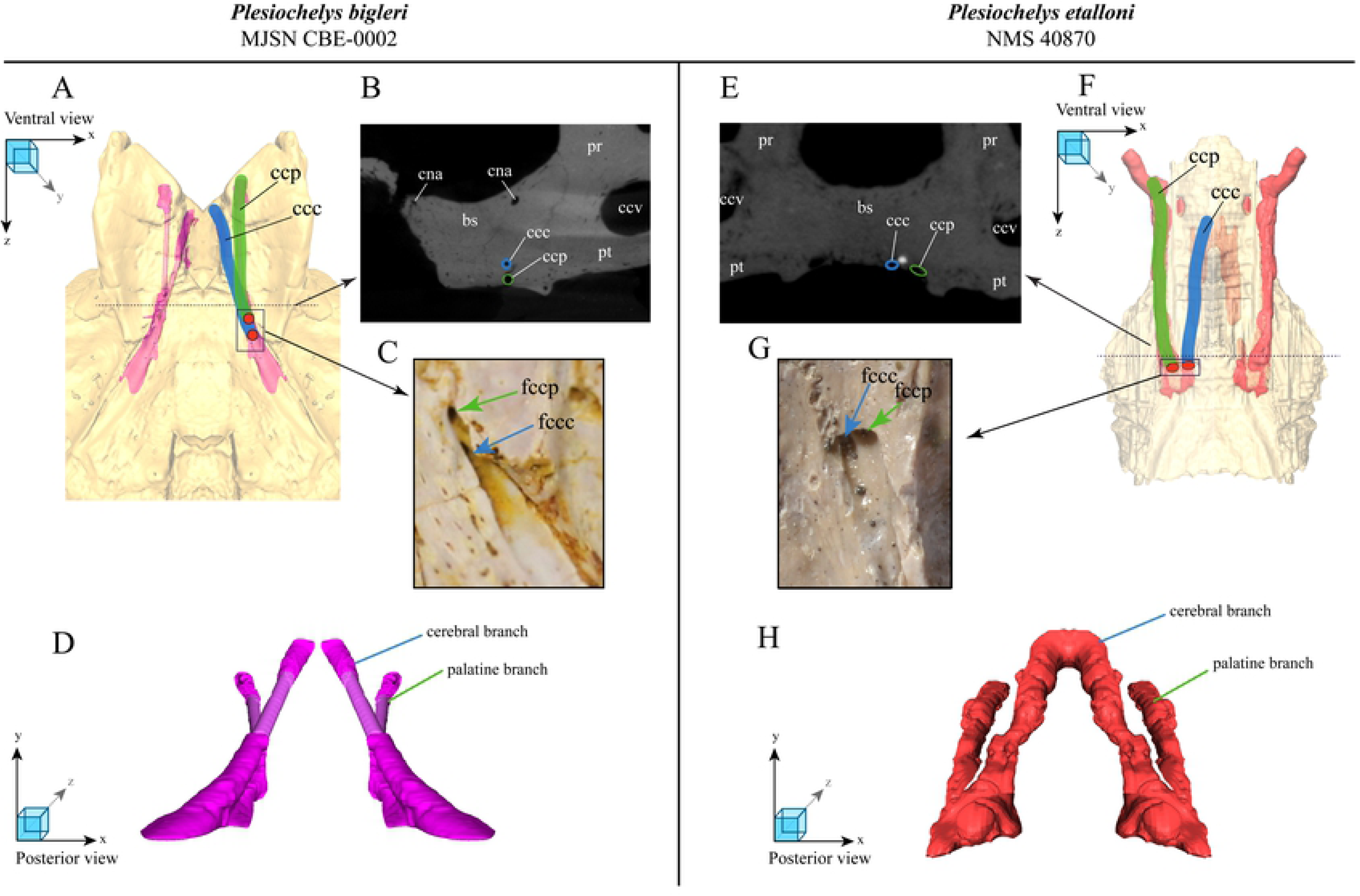
Spatial relation and course of the canalis caroticus cerebralis and canalis caroticus palatinum in MJSN CBE-0002 (A–D) and *Plesiochelys etalloni* (E–H). Spatial relation and course of the canalis caroticus cerebralis and canalis caroticus palatinum in MJSN CBE-0002 (A–D) and Plesiochelys etalloni (E-H). Schematic 3D model of the basisphenoid with the two branches of the carotid canal, in ventral view (the model is semitransparent and schematic - the more complete half was mirrored) for P.bigleri (A) and P.etalloni (F). Dashed line marks the position of the crosssection image. CT-cross section image (transversal plane) of the split between the cerebral and palatal branches in P. bigleri (B) and P. etalloni (E). Ventral view on the posterior foramina of the cerebral and palatine canal in P. bigleri (C) and P. etalloni (G). Schematic 3D model of the two branches of the carotid canal, in posterior view (the model is schematic - the more complete half was mirrored) for P.bigleri (D) and P.etalloni (H). Abbreviations: bs, basisphenoid; ccc, canalis carotici cerebralis; ccp, canalis carotici palatinum; cna, canalis nervi abducentis; fpccc, foramen posterior canalis carotici cerebralis; fpccp, foramen posterior canalis carotici palatinum.

The prootic is too poorly preserved to conclude on the presence or absence of an ossified pila prootica (an autapomorphy of *Plesiochelys etalloni*; [12,24]). On the dorsal surface of the otic chamber, there is a broad contact between the prootic and opisthotic, which prevents a contact between the supraoccipital and quadrate. This differs from most specimens referred to *Plesiochelys etalloni* [11]. As in the type material of *Plesiochelys bigleri*, the processus paroccipitalis extends posterolaterally. The foramen externum nervi glossopharyngei is also located quite laterally at the base of the processus interfenestralis. Due to the presence of matrix in this area, it is difficult to confirm the presence of a ridge extending anteriorly from the base of the processus interfenestralis, as in other specimens referred to *Plesiochelys bigleri*. MJSN CBE-0002 confirms the complete enclosure of the fenestra perilymphatica by bone in this taxon.

The basisphenoid is poorly preserved in MJSN CBE-0002, but what is visible, is consistent with the original description of *Plesiochelys bigleri* [11]. The anterior foramen nervi abducentis opens ventrally and slightly anteromedially to the base of the slightly damaged processus clinoideus, as in other plesiochelyids except *Plesiochelys etalloni*. The surface below the broken dorsum sellae appears to slope relatively gently anteriorly and presents a moderate midline tubercle. In the original description of *Plesiochelys bigleri*, Püntener et al. [11] stated that this surface is devoid of ridge or tubercle based on the morphology of the paratype specimen, but the cranium of the holotype specimen actually does have a moderate tubercle as in MJSN CBE-0002. The foramina anterius canalis carotici cerebralis are separated by a broad bar of bone and open slightly posterior to the level of the foramina anterius canalis carotici palatinum.

### Shell

MJSN CBE-0001 consists of several postcranial elements (see Material above), including an articulated fragment of carapace with the left costals 5–7 and part of the left peripherals 8 and 9 (Fig. 5). This fragment is 116 mm in length and 175 mm in width. MJSN CBE-0003 is a near complete shell measuring 450 mm in length and 405 mm in width (Fig. 6; data availability: surface scan of the shell). The plastron is 370 mm long and 320 mm wide. Superposing the articulated carapace fragment (MJSN CBE-0001) on top of the complete shell (MJSN CBE-0003) reveals that the two individuals were of the same size. MJSN CBE-0003 is one of the most complete shells referred to *Plesiochelys bigleri* and the best specimen to illustrate the three-dimensional shape of the shell in this taxon (see Discussion below).

**Fig. 5.**
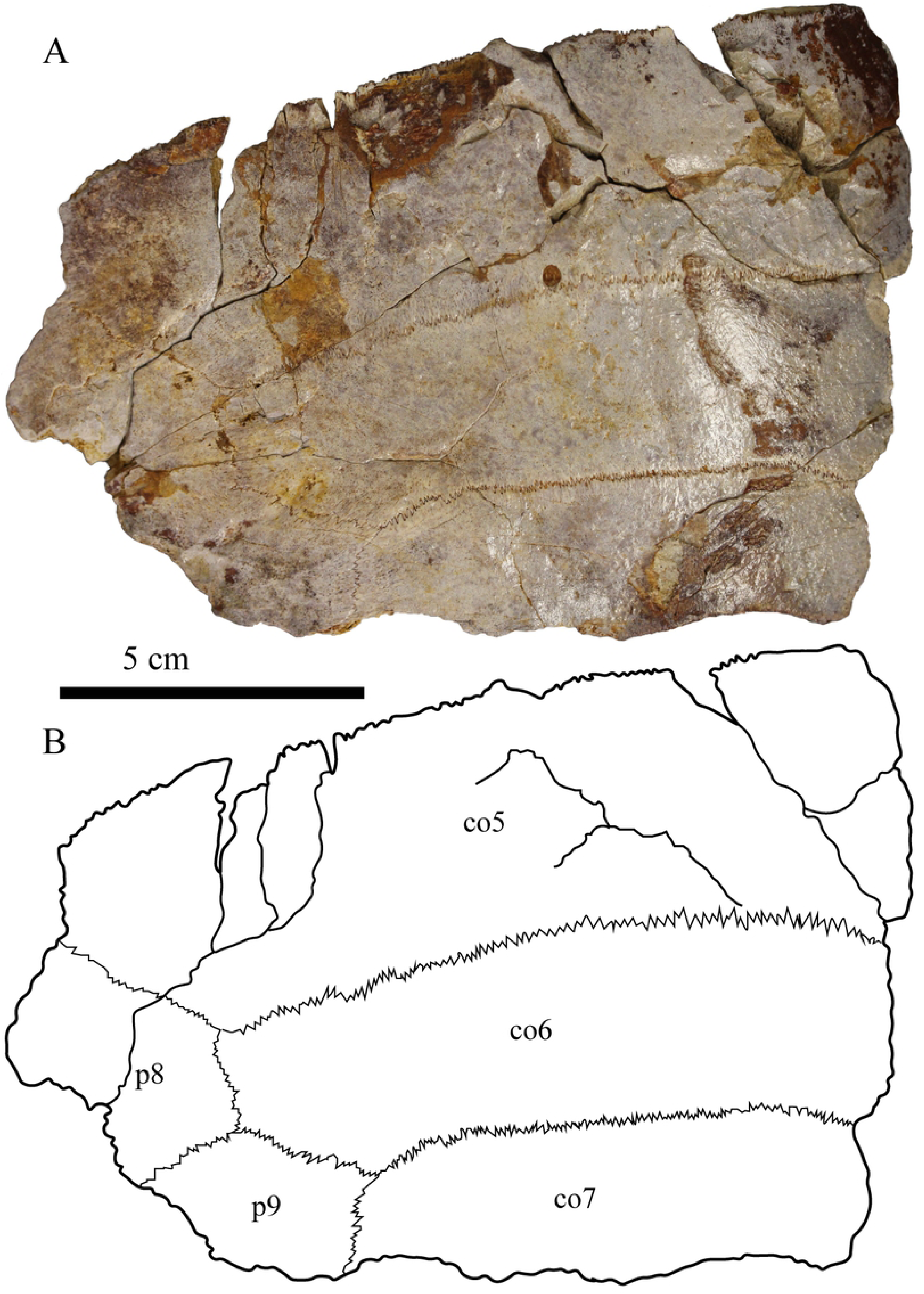
Carapace fragment MJSN CBE-0001 in dorsal view. Abbreviations: co, costals; p, peripherals.

**Fig. 6.**
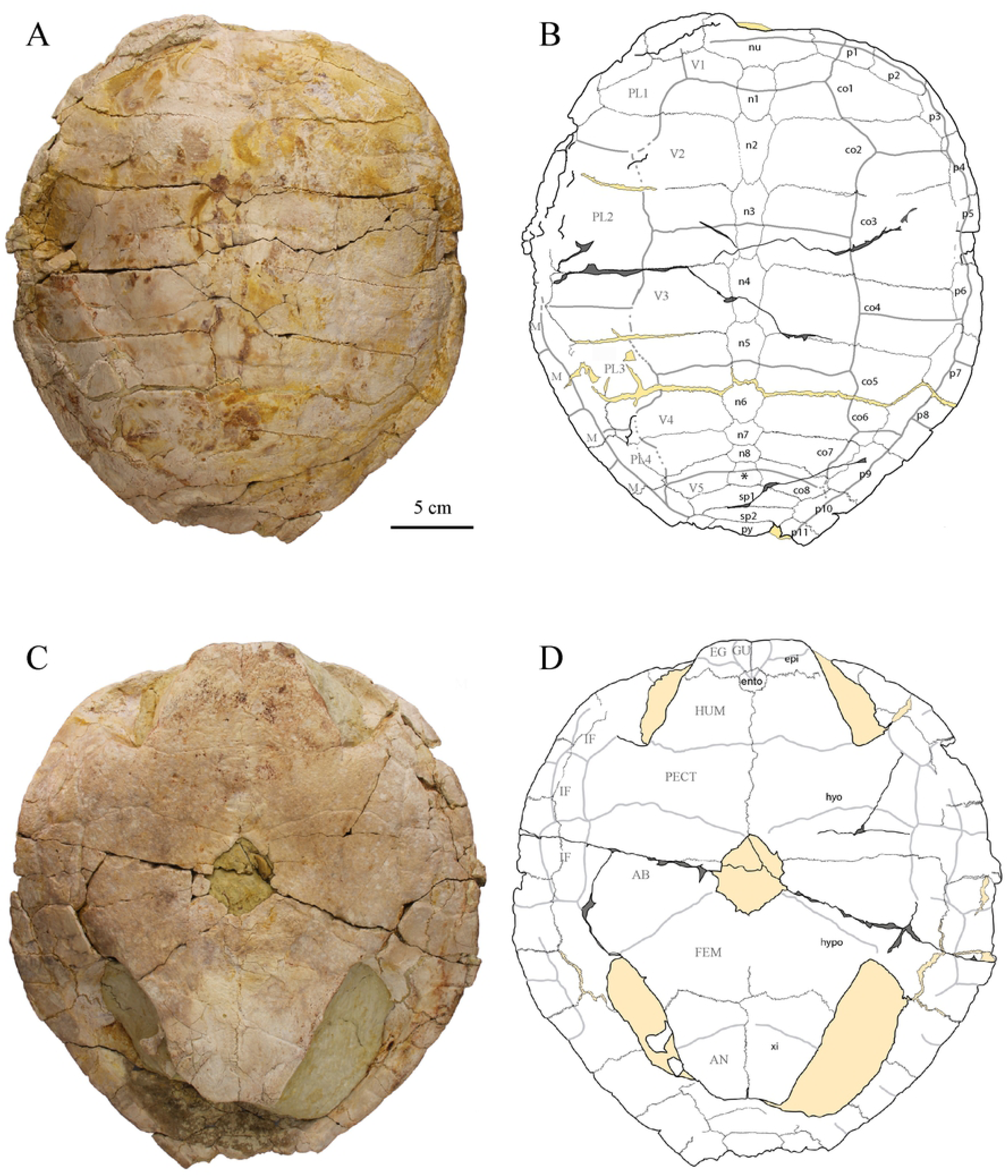
Shell MJSN CBE-0002. A-B: photograph and interpretative drawing of the carapace; C-D: photograph and interpretative drawing of the plastron. Abbreviations: AB, abdominal scale; AN, anal scale; co, costal; EG, extragular scale; ento, entoplastron; epi, epiplastron; FEM, femoral scale; GU, gular scale; HUM, humeral scale; hyo, hyoplastron; hypo, hypolastron; IF, inframarginal scale; M, marginal scale; n, neural; nu, nuchal; p, peripheral; PL, pleural scale; py, pygal; sp, suprapygal; V, vertebral scale; xi, xiphiplastron;*, intermediate element.

The outline of the carapace is roughly pentagonal with a moderately pointed posterior part (Fig. 6). The carapace is relatively low. Anteriorly, two parasagittal bulges, situated approximately at the medial third of the costals, frame a midline valley on the neurals (mostly visible at the level of neural 2 and costals 2). This part of the shell is not deformed, so this feature is probably natural. The general characteristics of the carapace of MJSN CBE-0003 agree fairly well with the description of Püntener et al. [11]: wider than long, trapezoidal nuchal, reduced nuchal notch, neural 1 oval in shape, neurals 2–6 elongated hexagons with shorter sides anteriorly, neurals 7 and 8 reduced in size (but preventing any midline contact between costals), roughly trapezoidal intermediate element tapering anteriorly, wide and trapezoidal suprapygals, eight pairs of costals and 11 pairs of peripherals, standard pattern of carapacial scutes (cervical region poorly preserved). The length-width ratio of costal 4 corresponds to what is known in *Plesiochelys etalloni* and other specimens referred to *Plesiochelys bigleri* (see [15]).

*Plesiochelys bigleri* and *Plesiochelys etalloni* can be differentiated based on neural bone thickness [11]. Unfortunately, this parameter cannot be precisely measured on the new material from Glovelier. Rough measurements were obtained by estimating the thickness of neurals 2–5 through fractures in or close to the neural elements with bands of paper. The resulting mean thickness and mean length/thickness ratio (Table 2; Fig. 7) are congruent with *Plesiochelys bigleri* and clearly out of the range for *Plesiochelys etalloni* (see [11]).

**Table 2:** Measurements of the neural elements of MJSN CBE-0003 and selected specimens of *Plesiochelys bigleri* and *Plesiochelys etalloni.* See Püntener et al. [11] for original data on *P. bigleri* and *P. etalloni*.

| Specimens | Mean neural 2–5<br>length [mm] | Mean neural 2–5<br>thickness [mm] | Length/thickness<br>ratio |
| --- | --- | --- | --- |
| MJSN CBE-0003 | 56.00 | 9.70* | 5.80 |
| *P. bigleri* (n=25) | 45.27–65.20 | 9.61–13.77 | 4.00–5.62 |
| *P. etalloni* (n=8) | 48.70–60.21 | 12.82–17.76 | 3.10–3.91 |

\*based on ten rough measurements (see text).

**Fig. 7.**
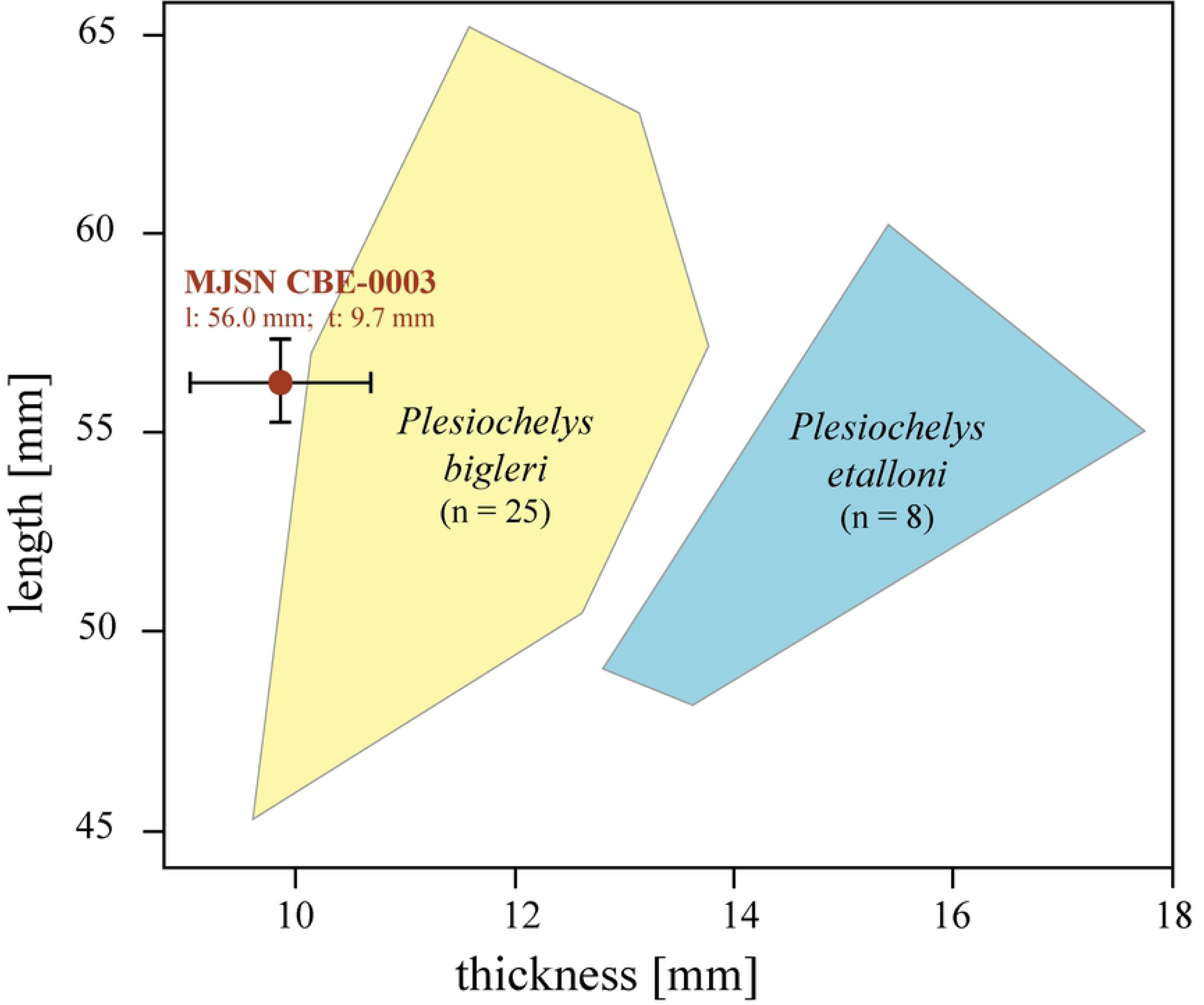
Mean neural length/thickness scatter plot for MJSN CBE-0002 (brown) and selected specimens of *Plesiochelys bigleri* (yellow) and *Plesiochelys etalloni* (blue). Graph modified after Püntener et al. [11]. Mean measurements of the new specimen MJSN CBE-0002 are plotted with the estimated range of error, which resulted from measurement limitations (see text).

The plastron of MJSN CBE-0003 presents an oval central plastral fontanelle. The central part of the plastron is moderately concave, which may indicate a male individual (by analogy with recent turtle species). The outline of the anterior plastral lobe is quadrangular, as typical in many specimens of *Plesiochelys bigleri* [11]. Epiplastral bulbs are absent. The entoplastron is rounded and rather small in comparison to most other known specimens of *Plesiochelys etalloni* and *Plesiochelys bigleri*, but an important variability in the shape and size of this element has been reported for these taxa [11,15]. The hyoplastron is very slightly wider than long. This should be noted, although other specimens referred to *Plesiochelys bigleri* have hyoplastra about as wide as long [11]. The xiphiplastra are rather long, as in many specimens of *Plesiochelys bigleri*, and noticeably angular posteriorly. The pattern of plastral scutes is conform to the original description [11].

### Postcranial elements

Of the right scapula (MJSN CBE-0001), only the dorsal scapular process is preserved. This process was crushed and broken in several places during fossilization (Fig. 8). As preserved, the scapular process is strongly arched and concave medially, but this is probably not its original shape. The scapular angle cannot be precisely measured, but can be estimated to be similar to what is known in *Plesiochelys* spp. [11,25].

**Fig. 8.**
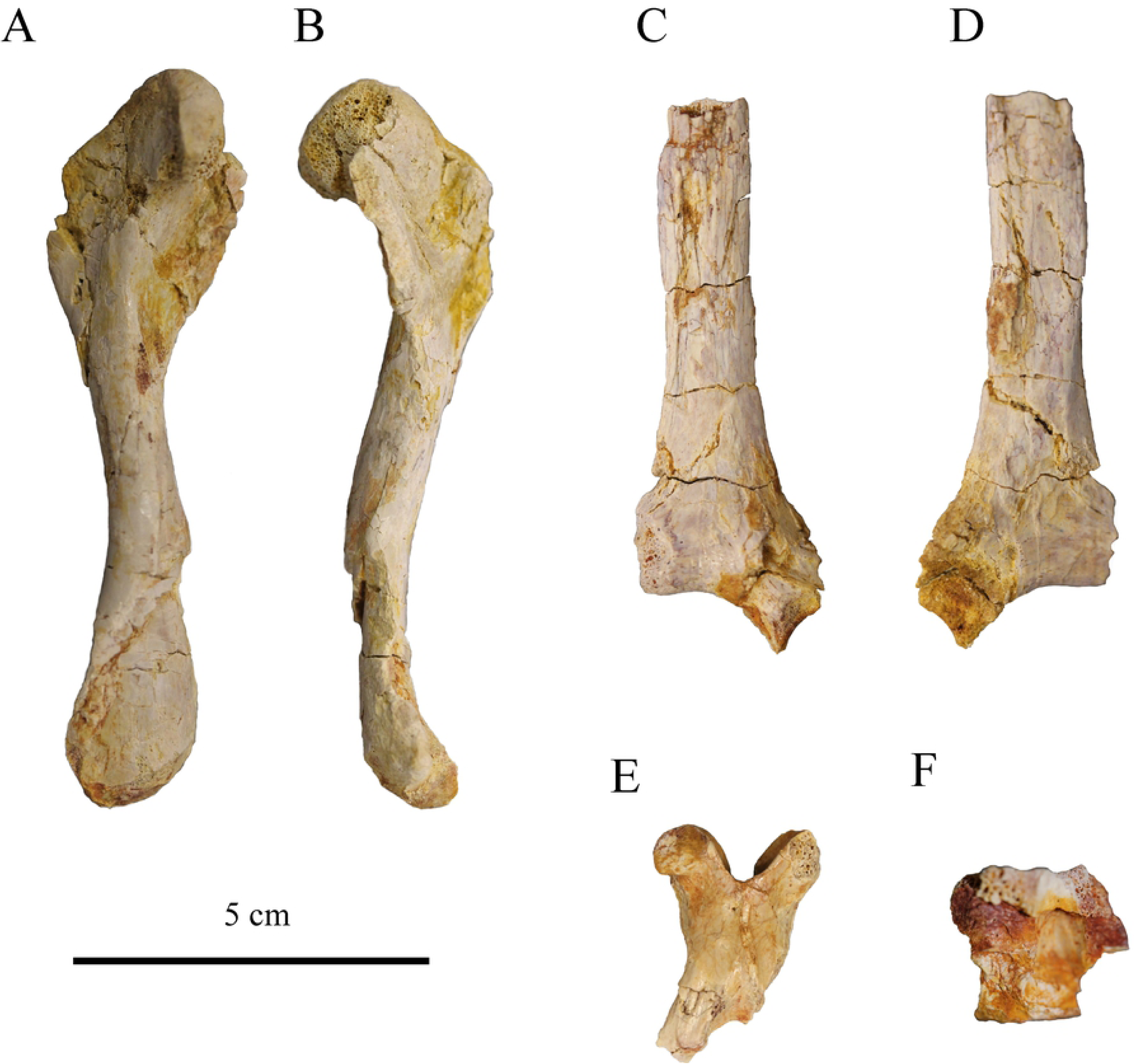
Postcranial material MJSN CBE-0001. Left femur in dorsal (A) and posterior (B) views; scapula in medial (C) and lateral (D) views; cervical vertebra in dorsal view (E); caudal vertebra in ventral view (F).

The femur is 100 mm long and relatively complete. Both trochanters and condyles are partly broken and the femoral head is damaged laterally. It was found in close association with the near complete shell MJSN CBE-0003. The morphology of the femur matches fairly well the description of Püntener et al. [11] and in that sense corresponds to what is known in other plesiochelyids, except *Tropidemys langii* [9].

A partial cervical vertebra was found associated with the near complete shell MJSN CBE-0003. It consists only of the neural arch, while the right pre- and postzygapophyses are missing. The posterior part of the neural arch is higher than the anterior part. The neural spine consists of a low longitudinal ridge. The pre-and postzygapophyses are widely spaced and their articular surface faces dorsomedially and ventrolaterally, respectively. This is consistent with what is known in other plesiochelyids and thalassemydids [11].

A small caudal vertebra centrum (12 mm long) was found associated with the shell MJSN CBE-0003. Interestingly, this centrum is amphicoelous with the two articular surfaces moderately concave. Bräm [25] described the caudal centra of *Plesiochelys etalloni* as procoelous, but this observation could not be reproduced recently [11]. The articular surfaces are oval in outline (slightly compressed dorsoventrally). There is a low, but robust ventral keel. A small nerve foramen is present just left of the ventral keel approximately midway along the centrum. The transverse process is apparently robust and located anteriorly along the centrum, but this part of the vertebra is poorly preserved.

## Discussion

### Alpha taxonomy

The new material from Glovelier can be confidently referred to *Plesiochelys bigleri* based on the following combination of characteristics: shallow pterygoid fossa, reduced processus trochlearis oticum, more rounded foramen nervi trigemini, anterior foramen nervi abducentis opening anteromedial to the base of the processus clinoideus, foramen anterius canalis carotici cerabralis located more anteriorly relative to the level of the dorsum sellae, superficial canalis caroticus internus (remaining possibly open ventrally), reduced posterior extension of the pterygoid over the cavum acustico-jugulare, processus paroccipitalis of the opisthotic extending posterolaterally, reduced neural bone thickness, epiplastral bulbs absent, and anterior plastral lobe quadrangular in outline (see [1,11]).

### New data on *Plesiochelys bigleri*

*Plesiochelys bigleri* was initially described based on a collection of 41 shells and two crania from the Kimmeridgian of the Porrentruy region, NW Switzerland [11]. The morphology of this taxon is therefore relatively well known. However, the new material from Glovelier described herein provided some additional information, which are summed up and discussed below.

The basicranium MJSN CBE-0002 provides unambiguous information on the contacts of the pterygoid with posterior basicranial elements. The pterygoid has a clear contact with the basioccipital, but lacks a contact with the exoccipital. The absence of a pterygoid-exoccipital contact is a difference with *Plesiochelys etalloni*, but the observation of this character is often difficult [12]. MJSN CBE-0002 also confirms the complete enclosure of the fenestra perilymphatica in *Plesiochelys bigleri*, but it is unclear whether the exoccipital or the basioccipital are responsible for the ventromedial closure of the fenestra. Finally, the new basicranium from Glovelier documents a new configuration of the split between the cerebral and palatine branches of the internal carotid artery (see below).

Although *Plesiochelys bigleri* is known by scores of shells, most were variably deformed and crushed during fossilization. A similar situation exists for shells of *Plesiochelys etalloni* from Solothurn and Porrentruy, the two most productive localities. However, the holotype of *P. etalloni* [14] and some specimens from Solothurn (e.g., NMS 8516, NMS 8579, NMS 8727, NMS 9173) suggest a low, evenly domed shell in this species. Most shells of *Plesiochelys bigleri* from the Porrentruy region were found in a semi-disarticulated state. Although attempts were made to estimate the 3D shape of the shell in this taxon by mounting the disarticulated elements on moldable sand [11], the reconstructions ranged from relatively flat to highly domed depending on the amount of postmortem deformation variably affecting the disarticulated shell bones. MJSN CBE-0003 therefore represents the first specimen to document the natural shell shape in *Plesiochelys bigleri*. The shell is relatively low, as in *P. etalloni*, but the two parasagittal bulges on the anterior costals are possibly a new character of *P. bigleri* (see data availability: surface scan of the shell).

Finally, the new material from Glovelier slightly extends the geographical and stratigraphical range of the species *Plesiochelys bigleri*. Most of the specimens known to date originate from the upper Kimmeridgian Lower *Virgula* Marls (Eudoxus ammonite zone), with only two incomplete shells found in the middle and upper parts of the lower Kimmeridgian Banné Marls (Cymodoce ammonite zone; see [11]). The new material from Glovelier was found close to the base of the Banné Marls and unambiguously confirms the occurrence of this species in this layer. This is also the first time the presence of the species is documented outside of the Porrentruy region.

### Carotid circulation

In most pan-cryptodires, the palatine branch of the internal carotid artery splits off laterally from the cerebral branch (e.g., [18,26]). This split usually occurs in the basisphenoid and pterygoid bones, and can therefore be documented in fossil turtles, though this may require special investigation tools (e.g., computed tomography). In some thalassochelydians, the canalis caroticus internus is superficial and is either unfloored or damaged so that the split between the cerebral and palatine branches of the internal carotid artery is apparent in ventral view. This is notably the case in most specimens referred to *Plesiochelys etalloni* [12]. In all specimens where the split is apparent externally, the palatine branch splits off laterally from the cerebral branch. Based on preliminary investigations, this is also the condition in the thalassochelydians *Jurassichelon oleronensis, Plesiochelys planiceps* and *Solnhofia parsonsi*. To our knowledge, MJSN CBE-0002 is unique among thalassochelydians in that the split between the cerebral and palatine branches occurs in a vertical plane. The palatine branch splits off ventrally from the carotid and travels ventral to the cerebral branch before entering the foramen posterius canalis carotici palatinum.

The skulls MJSN CBE-0002, MJSN TCH007-252 (holotype of *Plesiochelys bigleri*), and NMS 40870 (*Plesiochelys etalloni*) were CT-scanned and segmented for the purpose of this study (Supplementary material: S 1 & 2; data availability: isolated bones). Unfortunately, the canalis caroticus palatinum could not be reconstructed in MJSN TCH007-252. The paths of the canalis caroticus cerebralis and canalis caroticus palatinum were reconstructed for the two other crania (Fig. 4). In NMS 40870, the palatine branch splits off laterally and travels forward in the pterygoid while the cerebral branch continues dorsomedially through the basisphenoid. In MJSN CBE-0002, the palatine branch splits off ventrally and travels below the cerebral branch for a time. The two configurations are clearly different and confirm external observations.

As far as we know, the aforementioned configuration in MJSN CBE-0002 is unique in thalassochelydians, and possibly in pan-cryptodires, but further investigation is necessary. This feature can be added to the list of characteristics that differentiate *Plesiochelys bigleri* from *Plesiochelys etalloni*.

## Conclusion

The newly obtained material from Glovelier can be confidently referred to *Plesiochelys bigleri* and slightly extends the geographical and stratigraphical range of the species. The well-preserved cranium fragment (MJSN CBE-0002) reveals several morphological features and bone contacts that were not very clear or could not at all be observed in the type material of *Plesiochelys bigleri*. Especially noteworthy, is the spatial orientation of the split between the cerebral and palatine branches of the carotid artery. In thalassochelydians, this split occurs in a horizontal plane, while in this specimen, the split is oriented on a vertical plane instead. This condition seems to be unique for this clade. MJSN CBE-0003 is the first specimen to document the 3D shape of the shell of *Plesiochelys bigleri*. This rather low shell shows two parasagittal bulges on the anterior costals that may be considered as a new character of *P. bigleri.*

## Acknowledgements

We would like to thank Bernhard Hostettler (NMBE) for the help during the excavation of the material in 2014, for signaling the following material in 2016 and Ursula Menkveld–Gfeller (NMBE) for the access to the preparation lab at the Naturhistorische Museum in Bern. Likewise, we are grateful to Renaud Roch (MJSN) for the preparation of most of the material studied herein. Serjoscha Evers kindly provided access to segmented 3D models of the holotypes of *Plesiochelys planiceps* and *Solnhofia parsonsi*.

